# Prevalence of heritable symbionts in the Parisian Bedbugs (Hemiptera: Cimicidae)

**DOI:** 10.1101/2024.08.31.610657

**Authors:** Naciye Sena Cagatay, Mohammad Akhoundi, Arezki Izri, Sophie Brun, Gregory D. D. Hurst

## Abstract

Like many insects, the biology of bedbugs is impacted by a range of partner heritable microbes. Three maternally inherited symbionts are recognised, *Wolbachia* (an obligate partner), *Symbiopectobacterium* purcelli strain *Sy*Clec and *Ca. Tisiphia* sp. (facultative symbionts typically present in some but not all individuals). Past work had examined these heritable microbes from established laboratory lines, but not from broader field samples. We therefore deployed targeted endpoint PCR assays to determine the symbiont infection status for 50 bedbugs collected from 10 districts of Paris during the 2023 outbreak. All three symbionts were found to be broadly present across *C. lectularius* samples with the *Symbiopectobacterium*- *Ca*. Tisiphia-*Wolbachia* triple infection most commonly observed. A minority of individuals lacked either one or both facultative symbionts. Five mtDNA haplotypes were observed across the COI barcode region. Triple infections were found in all mtDNA haplotypes, indicating symbiont infection had been circulating over a protracted period of time. We conclude that the Parisian bedbug outbreak was one in which the host’s secondary symbionts were present at high frequency coinfections. Thus, facultative symbionts are an important but uncharacterized component of bedbug populations.

## Introduction

Hemipteran insects have diverse and important symbioses with microbial partners. Symbionts contribute to processes of digestion, anabolic activities (supply of vitamins & amino acids) (Akman et al., 2002), protection against natural enemies and defence against prey/hosts (Gosalbes et al., 2010), degradation of insecticides (Su et al., 2013), competence of host for transmitting viral diseases (Kliot and Ghanim, 2013). Parasitic interactions have known the maternal inheritance of the symbiont selecting for sex ratio distortion activity (Hurst and Frost, 2015).

The symbioses entered into by insects thus represent important drivers of insect biology. These interactions also offer novel routes for control of pest species and vector competence. For instance, obligate partners may be targetted as a means of killing focal parasites, as established for filarial nematodes where filariasis can be cured through targetted elimination of the worm’s *Wolbachia* symbionts (Simon et al. 1984)). Symbiont alteration of competence for viral transmission can also be effected through introduction of appropriate symbiont strains, and is widely used for control of zika and dengue transmission from mosquito hosts (Ant et al. 2023).

In this study, we examine the symbiotic associations of bedbugs. Bedbugs are a global pest whose blood-feeding activity has nuisance impacts (Dogget et al.,2012). Their nuisance status has particular reputational and financial impacts in the hospitality sector. Outbreaks of bedbugs – where infestations become locally frequency – are commonly observed and attain high press coverage. For instance, outbreaks of the common bedbug (*Cimex lecturalis* Linnaeus, 1758) and the tropical bedbug (*Cimex hemipterus* Fabricius, 1803) have occured regularly in Paris, France, with a notable peak event in hospitaility residences in 2023 (Chebbah et. al. 2023; Brimblecombe et al. 2024).

As obligate blood feeders, bedbugs development and reproduction rely on the presence of an obligate *Wolbachia* symbiont that supplies B vitamins to the host. Aposymbiotic bedbugs exhibit slower nymphal development, reduced adult survivorship, smaller adult size, fewer eggs per female, and a lower hatch rate comparing to bedbugs that harbor *Wolbachia* (Hickin et al. 2022, Kakumanu et al. 2024). Additionally, bed bugs experimentally depleted of *Wolbachia* can only develop with B vitamin supplementation, and the *Wolbachia* genome was found to contain a biotin synthesis operon within it (Hosokawa et al 2010; Nikoh et al 2014). Furthermore, *Wolbachia* is essential for various host reproductive processes such as cytoplasmic incompatibility, feminization, male death, parthenogenesis, increased or decreased fitness, and obligate symbiosis (Stouthamer et al. 1999, Gill et al. 2014, Nikoh et al. 2014. The *Wolbachia* found in bed bugs belongs to supergroup F (common in insects and filarial nematodes), with supergroup T recently reported in *C. hemipterus* from Senegal (Laidoudi et al. 2020).

Bedbugs additionally host two other heritable symbiont microbes, but these contrast with *Wolbachia* in not being required for the bedbug to develop and reproduce. They are therefore regarded as facultative symbionts, and indeed laboratory lines of bedbugs vary in the presence and frequency of these strains. First recognised was a gammaproteobacterium that falls in the recently described species *Symbiopectobacterium purcellii* (Nadal et al 2023). Described initially by Hypsa and Aksoy (1997) from a laboratory colony established in the USA, this microbe (henceforth *S. purcellii* strain *Sy*Clec) was later visualised as infecting the host ovaries alongside *Wolbachia* in Japanese bedbug strains (Hosokawa et al. 2010) and in the UK (Thongprem et al. 2021). More widely, *Symbiopectobacterium* is found associated with a variety of hemipteran insects, a member of the Hymenoptera and in a nematode (Tvedte et al. 2019; Martinson et al. 2020). Subsequently, a *Rickettsia* amplicon was detected in *C. lectularius* (Pilgrim et al. 2021). Additional reserach confirmed the maternal inheritance of this symbiont which was detected in 13 out of 21 laboratory strains from Europe and Africa (Thongprem et al. 2020). Infection with the symbiont was associated with a modest impact on host life history characters but with no evidence of reproductive parasitism (Thongprem et al. 2020). The symbiont has now been placed as *Ca. Tisiphia*, a genus sister to *Rickettsia* that includes many invertebrate heritable microbes (Davison et al. 2022).

Because the biology of insects is defined, in part, by the presence and frequency of symbionts, the presence and frequency of symbionts that are circulating are an important factor to consider in outbreaks. Previous research on bedbug heritable microbes has primarily focussed on diverse established laboratory lines, rather than bedbugs in the field context. We therefore analysed the pattern of presence and diversity of heritable symbionts in bedbugs collected across Parisian locations during the 2023 outbreak.

## Materials and methods

### Bed bug collection & identification

Bed bug samples were collected from various infested locations in the Paris area (Table 1). All the specimens were labeled and kept at -20°C for further molecular analysis.

**Table 1.** Collections sites of bedbugs in Paris in summer 2023.

| Location | Accommodation type |
| --- | --- |
| District 9 of Paris | Private apartment |
| District 11 of Paris | Private apartment |
| District 18 of Paris | Private apartment |
| District 19 of Paris | Private apartment |
| District 20 of Paris | Private apartment |
| Bobigny | HLM (social Collective apartment) |
| La Courneuve | Refugee camp residence |
| Pantin | Private apartment |
| Montreuil | Retirement home |
| Aubervilliers | HLM (social collective apartment) |

### DNA extraction,PCR amplification & sequencing

Bedbug DNA was extracted individually using Wizard® Genomic DNA Purification kit (Promega,USA). DNA was resuspended in 50 µl of molecular grade water and stored in −20°C for the future use. Amplification of the COI gene was initially assessed as a means of quality control by conventional PCR assay using the commonly used primer set (Thongprem et al. 2020; Chebbah et al. 2021). PCR assays consisted of a total of 15 μL per well, comprising 7.5 μL GoTaq® Hot Start Polymerase (Promega), 5.5 μL nuclease free water, 0.5 μL forward and reverse primers (concentration 10 pmol/μL) (Table S1) and 1 μL DNA template. COI barcode sequences (35 total) were additionally used to identify whether the respective bedbug specimen was *C. lectularius* or *C. hemipterus*. Where COI sequence was not obtained, an RFLP assay was conducted (digestion with *MspI* in 10x buffer Tango Thermoscientific), which cuts the COI amplicon of *C. hemipterus* but not *C. lecturalius*.

DNA template passing this QC stage were then utilized for onward detection of symbionts by PCR assay. Endpoint PCR targeting *Wolbachia* wsp gene, *Ca. Tisiphia* 16S rRNA & the citrate synthase (gltA) genes, and *Symbiopectobacterium* Gyrase B (GyrB) gene (Table S1: https://figshare.com/s/28c4ac95d12fdd361159). All assays used the following PCR conditions: initial denaturation at 95 °C for 5 min, followed by 35 cycles of denaturation (94°C for 30s), annealing (Ta°C for 30s), extension (72°C for 50s), and a final extension at 72°C for 7 min. The annealing temperature varied according to the primers (See Table S1 for primer sequences, Ta and expected product size). All PCR assays included positive (known infected) controls alongside blank (no DNA) negative controls.

PCR products were visualised on 1.5 % agarose gels stained with Midori Green Nucleic Acid Staining Solution (Nippon Genetics 41 Europe). From these data, each individual bedbug was allocated to species, and symbiont infection status (present/absent) for each of *Wolbachia, Symbiopectobacterium* and *Ca*. Tisiphia. In addition, mtDNA sequence of the individual was obtained in 35 cases.

Where positive amplication was observed for symbiont infection, the identity of the infection was ascertained by Sanger sequencing for one sample per location per symbiont using the forward primer. Sequences obtained were manutally curated and compared to known symbiont sequence for bedbugs by reference to Blastn homology Database (https://blast.ncbi.nlm.nih.gov/Blast.cgi).

### Phylogenetic Analyses

The resulting sequence chromatograms were edited and trimmed using Finch TV version 1.4.0 (© 2004–2006 Geospiza Inc.). Obtained sequences were submitted to GenBank under accession numbers PP887590-XXXXXX-XXXXXX.

The edited sequences were compared against known *Rickettsia* strains in the NCBI database. *Occidentia massiliensis* was selected as the outgroup for phylogenetic analysis of both genes (16S rRNA & gltA). All selected *Rickettsia* sequences were then aligned with the new sequences using MAFFT v7.4 (Katoh et al. 2002). ModelFinder (Kalyaanamoorthy et al., 2017) was employed to select the best-fit models. Maximum likelihood phylogenetic trees were then inferred for both genes using IQ-TREE (Nguyen et al., 2015) with 1,000 rapid ultra fast bootstrap replicates (Minh et al., 2020). TN+F+I and HKY+F+G4 models were used for the 16S rRNA and gltA genes, respectively.

*Symbiopectobacterium* gyrase B dataset was aligned with MAFFTv7.4 (Katoh et al. 2002). Modelfinder was used to ascertain the appropriate model (TIM2e+G4). Phylogenetic relatedness was estimated using IQTREE (Nguyen et al., 2015) using 1,000 rapid bootstrap replicates (Minh et al., 2020).

### Concordance of bed bugs mtDNA sequences and their symbiont infection status

The COI sequence for *C. lectularius* above were aligned with MAFFTv7.4 (Katoh et al. 2002). Haplotype diversity indices were calculated for bed bug populations in DnaSP v6.12.03 (Rozas et al., 2003). Subsequently, the haplotypes were extracted, and a haplotype-spanning network was estimated using the TCS haplotype network algorithm generated in PopART v7 (Leigh et al. 2015). Symbiont infection status was then mapped to mtDNA haplotype.

### Statistical Analyses

The statistical analyses were performed using *C*.*lecturalis* samples collected from various locations. The data were analyzed using the statistical platform R (version 4.3) (R Development Core Team, 2022) with the packages “Companion to applied regression(car)” (Fox and Weisberg, 2019). To explore associations between endosymbionts, double infection status, and the mtDNA haplotype groups and *C. lecturalis* populations collected from various locations, a Fisher’s exact test was performed, with a significance cut-off set at *P* <0.005.

## Results

### Prevalence of endosymbionts bacteria results

Of the 50 bedbugs tested from 10 locations, 45 were *C. lectularius* and five were identified as *C. hemipterus*.

*Wolbachia* was detected by species-specific endpoint PCR (wsp gene) in all *C*.*lectularius* and *C. hemipterus* specimens. *Cimex hemipterus* was not observed to be infected with either of the facultative symbionts. The facultuative symbionts *Symbiopectobacterium* and *Ca. Tisiphia* were detected in *C. lectularius* (Table 2; https://figshare.com/s/96ce0e00318c1caaae11). Infection with these symbionts occured broadly over space, and individuals carrying all three symbionts were most common. Tests revealed no evidence to reject the null hypothesis that *Symbiopectobacterium* infection frequency occured evenly between locations (Fisher’s exact test, *P>* 0.05). In contrast, heterogeneity between samples was supported for *Ca. Tisiphia* infection, where some samples had all individuals infected, and some no infected individuals (Fisher’s exact test, *P* <0.05).

**Table 2.** Infection status of *C. lectularius* individuals from various areas of Paris. Sy+ = positive for *Symbiopectobacterium*, Sy-negative or *Symbiopectobacterium*; T+ = positive for *Ca*.*Tisiphia*, T-negative for *Ca. Tisiphia*

| Location | Sy+ T+ | Sy+ T- | Sy-T+ | Sy-T- |
| --- | --- | --- | --- | --- |
| District 9 of Paris | 5 | 0 | 0 | 0 |
| District 11 of Paris | 2 | 2 | 1 | 0 |
| District 18 of Paris | 0 | 1 | 0 | 0 |
| District 19 of Paris | 4 | 1 | 0 | 0 |
| District 20 of Paris | 0 | 4 | 0 | 1 |
| Bobigny | 5 | 0 | 0 | 0 |
| La Courneuve | 0 | 5 | 0 | 0 |
| Pantin | 2 | 1 | 1 | 1 |
| Montreuil | 5 | 0 | 0 | 0 |
| Aubervilliers | 2 | 0 | 1 | 1 |
| **Total Paris** | **25** | **14** | **3** | **3** |

Amplicon sequences obtained for Ca. *Tisiphia* (*gltA*, 16S rRNA) and *Symbiopectobacterium* (*gyrB*) were the same across all individuals examined. The sequences obtained were aligned to *Symbiopectobacterium* sequences previously described in this species. Phylogenetic analysis compared to other known strains of symbiont clearly clustered the Parisian isolates as monophyletic with previous recorded strains from bedbugs (Figure 1). The *Wolbachia* sequence from our *C. lectularius* matched 100% with the *Wolbachia* endosymbiont of *C. lectularius* previously reported in this species (KR706518.1).

**Figure 1.**
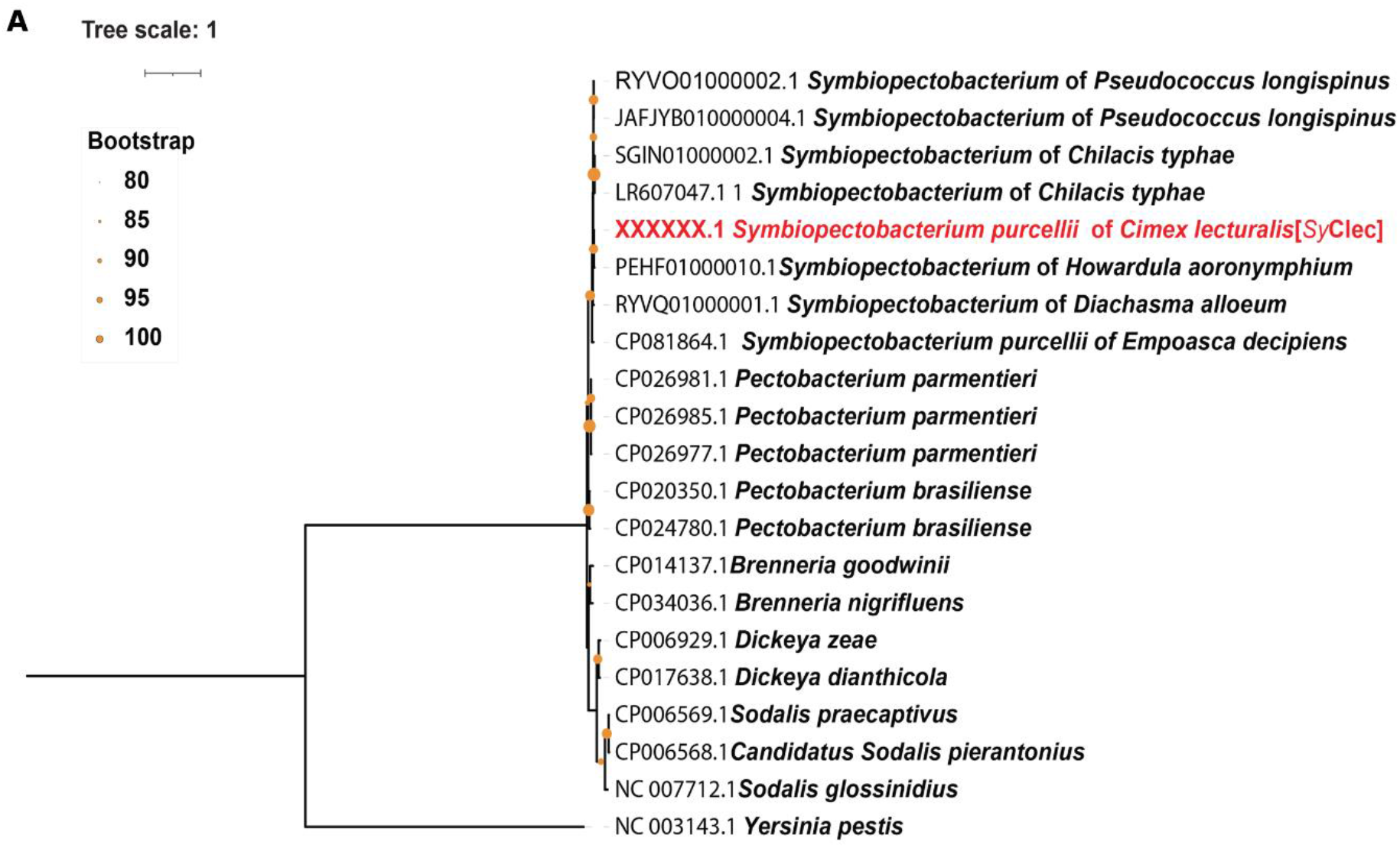
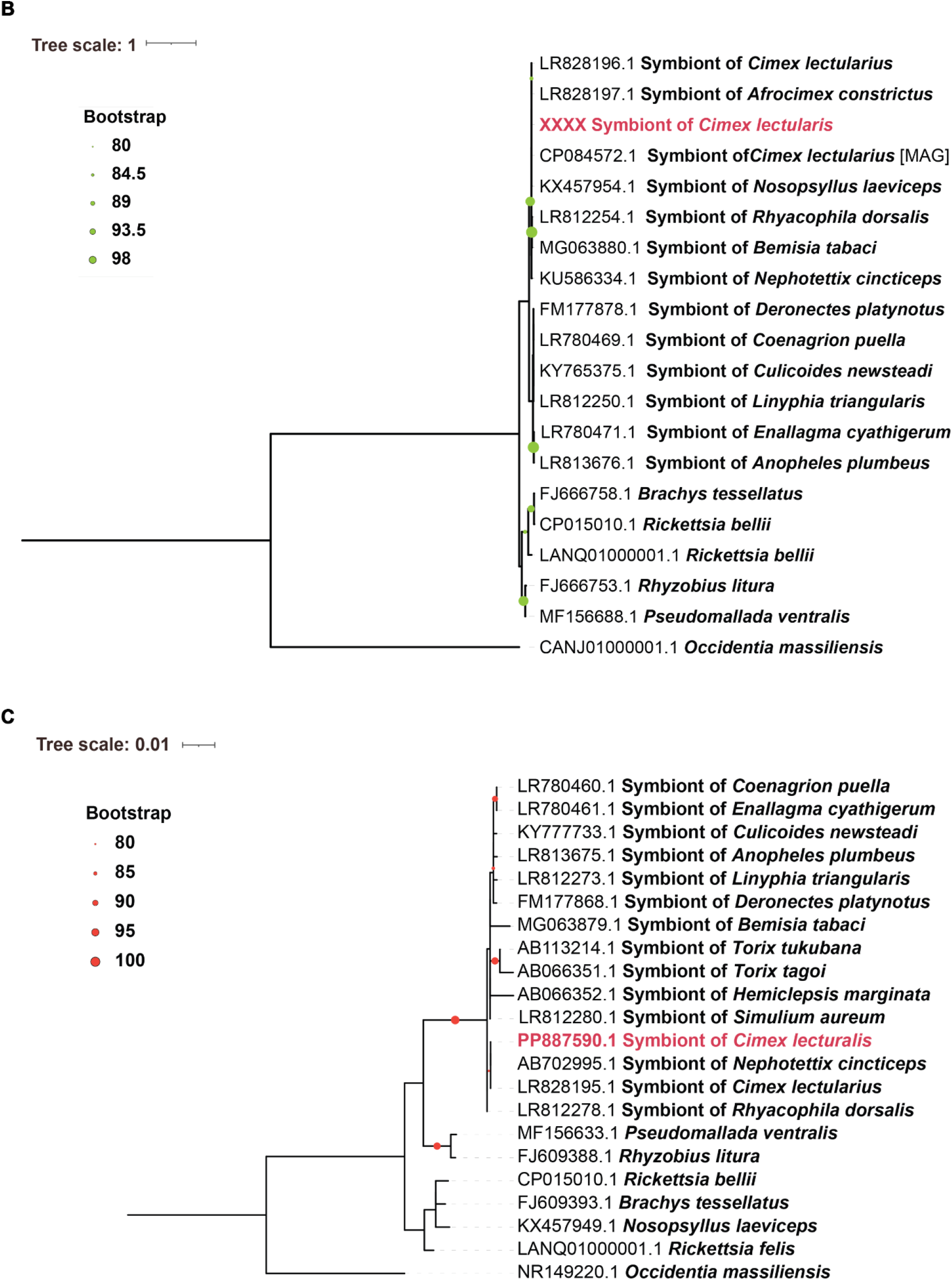
Maximum likelihood phylogenetic trees of facultative symbionts of Parisian *C. lectularius* constructed using gyrB of *Symbiopectobacterium* (A), *gltA*,of *Ca. tisiphia* (B) and 16S rRNA of *Ca. tisiphia* (C) gene sequences compared to previously recorded congeneric symbiont sequences originating from GenBank names given correspond to the host species of the symbiont.

We then mapped symbiont infection status onto mtDNA haplotype network (Figure 2). This analysis revealed five mtDNA haplotypes defined by three SNPs across 257 bp of high quality score sequence (>30). The *Symbiopectobacterium/ Ca*. Tisiphia coinfection was found in all five mtDNA haplotype obtained; individuals with a single infection (Sy+T-, Sy-T+) or no facultative symbiont infection (Sy-T-) were found in a subset of mtDNA haplotypes.

**Figure 2.**
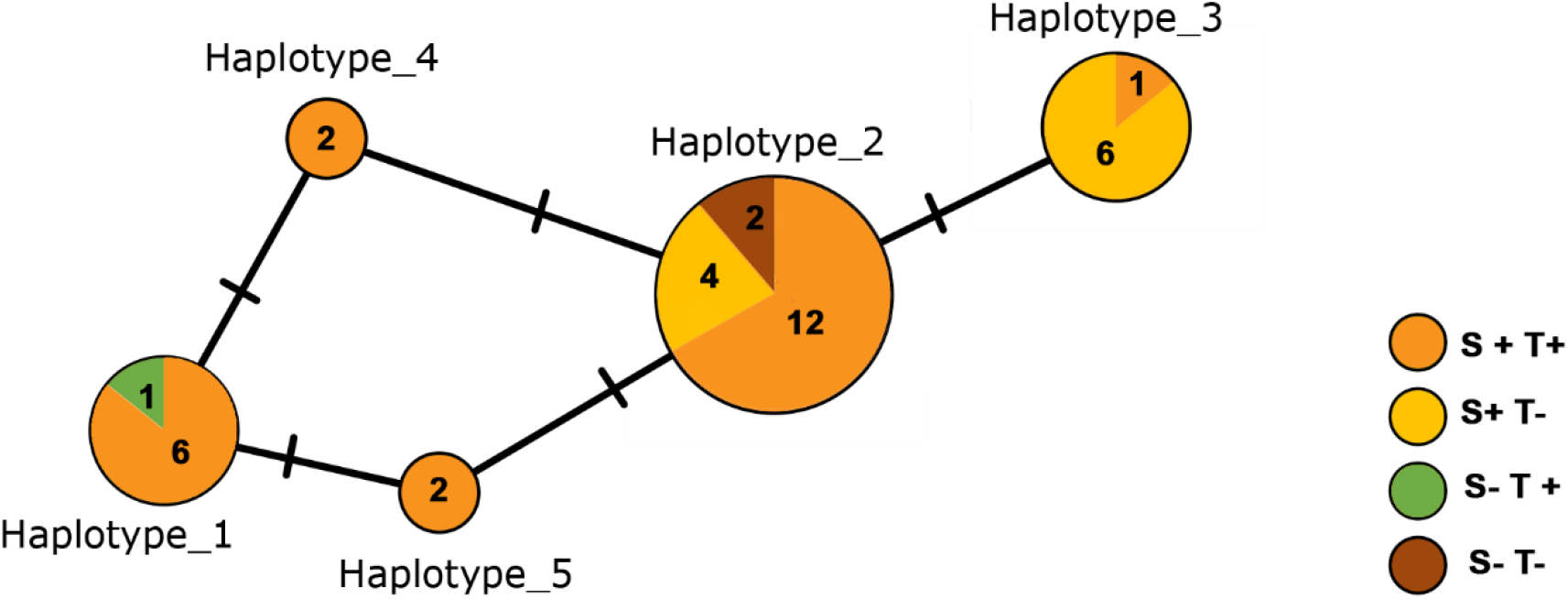
Haplotype spanning network for mtDNA in Parisian *C. lectularius* constructed from 257bp of COI sequences, partitioned by symbiont infection status. Size of circle indicates number of individuals with that haplotype; colours within represent facultative symbiont infection statust of the individuals with that haplotype (Sy+= positive for *Symbiopectobacterium*, Sy-negative for *Symbiopectobacterium*; T+ = positive for *Ca*.*Tisiphia*, T-negative for *Ca. Tisiphia*).

## Discussion

Bedbugs are amongst the most common ectoparasites of humans worldwide. While not regarded as biological vectors, they are a serious nuisance pest with a high economic impact. Like many insects, the biology of bedbugs is partly influenced by their symbiotic partners. For *C. lectularius*, these include the obligate symbiont *Wolbachia*, as well as the facultative symbionts *Symbiopectobacterium Sy*Clec and *Ca*. Tisiphia. The presence of facultative symbionts is largely documented in laboratory lineages, while our research focuses on exploring their diversity and frequency of infection in field-caught bed bug populations and demonstrates their circulaton at high frequency in field-collected samples. However, *Symbiopectobacterium Sy*Clec and *Ca*. Tisiphia were not present in all individuals sampled, corroborating its status as a facultative partner.

To this end, we examined the heritable microbiota of bedbugs collected during the Paris outbreak of summer 2023 using a targeted PCR approach. These data represent the first survey of field-collected bedbugs for the three symbionts. The predominant bedbug species recovered was *C. lectularius*, and all three previously described symbionts from this species were recovered in our samples. Indeed, the majority of individuals collected were coinfected with all three symbionts, *Wolbachia, Ca. Tisiphia* and *Symbiopectobacterium*. Other individuals carried either *Wolbachia* alone, or *Wolbachia* along with one of the other two symbiont strains. Theoretical work indicates that where coinfected forms are common, individuals lacking one or more of the infections are expected to be found due to segregational loss (inefficient maternal inheritance) (Frank,1998). The high frequency of triply infected individuals alongside a minority of bedbugs with either just one or neither facultative symbiont make it likely that the triply infected form occasionally shows segregation of the facultative symbionts, creating doubly infected individuals, and these doubly infected individuals occasionally show segregation of the remaining facultative symbiont resulting in *Wolbachia*-only infected lines. This model is consistent with data from maintained laboratory lines, where particular isofemale matrilines have a mix of *Ca. Tisiphia* infected and uninfected individuals (Thongprem et al. 2020).

In addition, we examined the relationship between symbiont status and mitochondrial variation. Mitochondria are also maternally inherited, such that the dynamics of symbionts impact that of mitochondria and *vice versa* (Galtier et al 2009; Fenton et al 2021). This data recovered five mtDNA haplotypes from the COI amplicon sequence. Triply infected bedbugs were represented in all five haplotypes identified, and doubly/singly infected individuals in a subset.The diversity of mtDNA types found indicates there has not been a recent selective sweep of either mtDNA or symbionts, as recent selective sweeps of symbionts or mtDNA are associated with absence of mtDNA variation. Our data, in contrast, indicates that the symbionts have likely been circulating in the species for some time. This conclusion is consistent with *Symbiopectobacterium Sy*Clec and *Ca*. Tisiphia being recorded from *C. lectularius* across continents in previous work (Hypša and Aksoy,1997; Thongpremet al. 2020; Pilgrim et. al. 2021; Davison et al. 2022). Further, the presence of doubly and singly infected individuals in only a subset of mtDNA haplotypes is consistent with their creation by segregation of symbionts from triply and doubly infected individuals as described above.

Past work has emphasized the functional significance of *Wolbachia* to bedbugs, and indicated the presence of *Ca. Tisiphia* and *Symbiopectobacterium Sy*Clec. Our study provides population-level data for the frequency of these facultative partners in the Parisian outbreak, and indicates that triply infected individuals are the most common. Furthermore, the mtDNA data indicates that the symbionts have been present for a sufficient evolutionary time that they are associated with diverse mtDNA haplotypes. What is not clear is their functional significance. Their persistence over time and their circulation at high frequency, combined with evidence that they do not cause reproductive parasitism (Thongprem et al. 2020), make it most parsimonious to consider them as partners of unknown (but significant) benefit.

## Supporting information

https://figshare.com/s/28c4ac95d12fdd361159

## Acknowledgements

N.S.C. was supported by TÜBİTAK [The Scientific and Technological Research Council of Türkiye, project number: 1059B192202296] 2219-International Postdoctoral Research Fellowship Program.

## Data availability

The raw data underpinning Table S1(https://figshare.com/s/28c4ac95d12fdd361159) and raw data(https://figshare.com/s/96ce0e00318c1caaae11) can be accessed at figshare. Marker gene sequences are available under accessions xxxxxx-PP886873-PP886877.

## CReDIT roles

Conceptualization: NSC, MA, GH

Data curation: NSC

Formal Analysis: NSC, GH

Funding acquisition: NSC

Investigation: NSC, MA, AI, SB

Methodology: NSC, GH

Project administration: GH, MA

Resources: GH, MA

Supervision: GH, MA

Validation: NSC

Visualization: NSC

Writing – original draft: SC, MA, GH

Writing – review & editing: all authors.

## References

Akman, L., Yamashita, A., Watanabe, H., Oshima, K., Shiba, T., Hattori, M. and Aksoy, S., 2002. Genome sequence of the endocellular obligate symbiont of tsetse flies, Wigglesworthia glossinidia. Nature genetics, 32(3), pp.402–407.

Ant, T.H., Mancini, M.V., McNamara, C.J., Rainey, S.M. and Sinkins, S.P., 2023. Wolbachia-Virus interactions and arbovirus control through population replacement in mosquitoes. Pathogens and Global Health, 117(3), pp.245–258.

Brimblecombe, P., Mueller, G. and Querner, P., 2024. Public and media interest in bed bugs-Europe 2023. Current Research in Insect Science, 5, p.100079.

Chebbah, D., Elissa, N., Sereno, D., Hamarsheh, O., Marteau, A., Jan, J., Izri, A. and Akhoundi, M., 2021. Bed bugs (Hemiptera: Cimicidae) population diversity and first record of Cimex hemipterus in Paris. Insects, 12(7), p.578.

Chebbah, D., Hamarsheh, O., Sereno, D., Elissa, N., Brun, S., Jan, J., Izri, A. and Akhoundi, M., 2023. Molecular characterization and genetic diversity of Wolbachia endosymbionts in bed bugs (Hemiptera; Cimicidae) collected in Paris. Plos one, 18(9), p.e0292229.

Davison, H.R., Pilgrim, J., Wybouw, N., Parker, J., Pirro, S., Hunter-Barnett, S., Campbell, P.M., Blow, F., Darby, A.C., Hurst, G.D. and Siozios, S., 2022. Genomic diversity across the Rickettsia and ‘Candidatus Megaira’genera and proposal of genus status for the Torix group. Nature Communications, 13(1), p.2630.

Doggett, S. L., Dwyer, D. E., Peñas, P. F., & Russell, R. C. (2012). Bed bugs: clinical relevance and control options. Clinical microbiology reviews, 25(1), 164–192.

Fenton, A., Camus, M.F. and Hurst, G.D., 2021. Positive selection on mitochondria may eliminate heritable microbes from arthropod populations. Proceedings of the Royal Society B, 288(1959), p.20211735.

Frank, S.A., 1998. Foundations of social evolution (Vol. 19). Princeton University Press.

Fox, J. and Weisberg, S., 2019. Nonlinear regression, nonlinear least squares, and nonlinear mixed models in R. population, 150, p.200.

Galtier, N., Nabholz, B., Glémin, S. and Hurst, G.D.D., 2009. Mitochondrial DNA as a marker of molecular diversity: a reappraisal. Molecular ecology, 18(22), pp.4541–4550.

Gosalbes, M.J., Latorre, A., Lamelas, A. and Moya, A., 2010. Genomics of intracellular symbionts in insects. International Journal of Medical Microbiology, 300(5), pp.271–278.

Hickin ML, Kakumanu ML, Schal C. Effects of Wolbachia elimination and B-vitamin supplementation on bed bug development and reproduction. Sci Rep. 2022 Jun 17;12(1):10270.

Hosokawa, T., Koga, R., Kikuchi, Y., Meng, X.Y. and Fukatsu, T., 2010. Wolbachia as a bacteriocyte-associated nutritional mutualist. Proceedings of the National Academy of Sciences, 107(2), pp.769–774.

Hurst, G.D. and Frost, C.L., 2015. Reproductive parasitism: maternally inherited symbionts in a biparental world. Cold Spring Harbor perspectives in biology, 7(5), p.a017699.

Hypša, V. and Aksoy, S., 1997. Phylogenetic characterization of two transovarially transmitted endosymbionts of the bedbug Cimex lectularius (Heteroptera: Cimicidae). Insect molecular biology, 6(3), pp.301–304.

Kakumanu ML, Hickin ML, Schal C. Detection, Quantification, and Elimination of Wolbachia in Bed Bugs. Methods Mol Biol. 2024;2739:97–114.

Kalyaanamoorthy, S., Minh, B.Q., Wong, T.K., Von Haeseler, A. and Jermiin, L.S., 2017. ModelFinder: fast model selection for accurate phylogenetic estimates. Nature methods, 14(6), pp.587–589.

Katoh, K., Misawa, K., Kuma, K.I. and Miyata, T., 2002. MAFFT: a novel method for rapid multiple sequence alignment based on fast Fourier transform. Nucleic acids research, 30(14), pp.3059–3066.

Kliot, A., Cilia, M., Czosnek, H. and Ghanim, M., 2014. Implication of the bacterial endosymbiont Rickettsia spp. in interactions of the whitefly Bemisia tabaci with tomato yellow leaf curl virus. Journal of virology, 88(10), pp.5652–5660.

Laidoudi, Y., Levasseur, A., Medkour, H., Maaloum, M., Ben Khedher, M., Sambou, M., Bassene, H., Davoust, B., Fenollar, F., Raoult, D. and Mediannikov, O., 2020. An earliest endosymbiont, Wolbachia massiliensis sp. nov., strain PL13 from the bed bug (Cimex hemipterus), type strain of a new supergroup T. International journal of molecular sciences, 21(21), p.8064.

Leigh, J.W., Bryant, D. and Nakagawa, S., 2015. POPART: full-feature software for haplotype network construction. Methods in Ecology & Evolution, 6(9).

Leslie, J.F., 1984. A “sex-ratio” condition in Oncopeltus fasciatus. Journal of Heredity, 75(4), pp.260–264.

Martinson, V.G., Gawryluk, R.M., Gowen, B.E., Curtis, C.I., Jaenike, J. and Perlman, S.J., 2020. Multiple origins of obligate nematode and insect symbionts by a clade of bacteria closely related to plant pathogens. Proceedings of the National Academy of Sciences, 117(50), pp.31979–31986.

Minh BQ, Schmidt HA, Chernomor O et al. IQ-TREE 2: New models and efficient methods for phylogenetic inference in the genomic era. Molecular Biology and Evolution 2020;37:1530–4. 10.1093/molbev/msaa015

Nadal-Jimenez, P., Siozios, S., Halliday, N., Cámara, M. and Hurst, G.D., 2022. Symbiopectobacterium purcellii, gen. nov., sp. nov., isolated from the leafhopper Empoasca decipiens. International Journal of Systematic and Evolutionary Microbiology, 72(6), p.005440.

Nguyen, L.T., Schmidt, H.A., Von Haeseler, A. and Minh, B.Q., 2015. IQ-TREE: a fast and effective stochastic algorithm for estimating maximum-likelihood phylogenies. Molecular biology and evolution, 32(1), pp.268–274.

Nikoh, N., Hosokawa, T., Moriyama, M., Oshima, K., Hattori, M. and Fukatsu, T., 2014. Evolutionary origin of insect–Wolbachia nutritional mutualism. Proceedings of the National Academy of Sciences, 111(28), pp.10257–10262.

Pilgrim, J., Thongprem, P., Davison, H.R., Siozios, S., Baylis, M., Zakharov, E.V., Ratnasingham, S., DeWaard, J.R., Macadam, C.R., Smith, M.A. and Hurst, G.D., 2021. Torix Rickettsia are widespread in arthropods and reflect a neglected symbiosis. GigaScience, 10(3), p.giab021.

R Development Core Team. (2022). R: a language and environment for statistical computing. R Foundation for Statistical Computing, Vienna, Austria. ISBN 3-900051-07-0, http://www.R-project.org

Rozas, J., Sánchez-DelBarrio, J.C., Messeguer, X. and Rozas, R., 2003. DnaSP, DNA polymorphism analyses by the coalescent and other methods. Bioinformatics, 19(18), pp.2496–2497.

Su, Q., Zhou, X. and Zhang, Y., 2013. Symbiont-mediated functions in insect hosts. Communicative & integrative biology, 6(3), p.e23804.

Thongprem, P., Evison, S.E., Hurst, G.D. and Otti, O., 2020. Transmission, tropism, and biological impacts of torix rickettsia in the common bed bug Cimex lectularius (Hemiptera: Cimicidae). Frontiers in microbiology, 11, p.608763.

Tvedte, E.S., Walden, K.K., McElroy, K.E., Werren, J.H., Forbes, A.A., Hood, G.R., Logsdon Jr, J.M., Feder, J.L. and Robertson, H.M., 2019. Genome of the parasitoid wasp Diachasma alloeum, an emerging model for ecological speciation and transitions to asexual reproduction. Genome biology and evolution, 11(10), pp.2767–2773.

